# A fluorescent reporter system for scalable and live detection of PYY production from enteroendocrine cells with single-event resolution

**DOI:** 10.64898/2026.04.24.720392

**Authors:** Aanya Hirdaramani, Gary Frost, Aylin C. Hanyaloglu

## Abstract

Peptide YY (PYY) comprises the secretory repertoire of enteroendocrine L-cells alongside glucagon-like peptide-1 (GLP-1), and positively modulates postprandial satiety, digestion mechanics, and regeneration of the intestinal epithelium. Whereas immortalised GLP-1-secreting human L-cell models support pre-clinical drug discovery, comparable human lines that robustly secrete PYY are lacking, hindering mechanistic studies of its release. We present a biosensor for scalable detection of PYY production and secretion from human enteroendocrine cells *in vitro.* Guided by *in silico* structural prediction, we engineer a superecliptic phluorin (SEP)-tagged PYY, SEP-PYY, that engages native hormone processing machinery and is responsive to canonical nutrient stimuli when expressed in a human enteroendocrine cell line, NCI-H716. SEP-PYY production and secretion can be measured by optimised flow cytometry and spectrofluorometric plate readouts respectively, methods with superior time- and cost-efficacy to current hormone detection methods. Leveraging the pH sensitivity of SEP, we use this reporter system in detection of single-event hormone exocytosis by total internal reflection microscopy. Finally, we demonstrate the application of our system in screening ligands of metabolite-sensing G-protein coupled receptors that drive SEP-PYY secretion, and supporting discovery of druggable pathways in metabolic disease.

## Main

Peptide YY (PYY) is secreted by enteroendocrine L-cell populations enriched in the distal gut, in response to nutrient and microbiota-derived stimuli^1^. Acting through neuropeptide Y (NPY) receptors, PYY regulates central appetite pathways^2^, modulates gastrointestinal motility and nutrient absorption^3^, and exerts growth factor–like trophic effects on epithelial and endocrine tissues. The anorexigenic effects of PYY are primarily mediated by Y2R^4,5^, and Y2R agonism has demonstrated marked pre-clinical efficacy in reduction of weight loss and energy intake when administered as a co-therapy to GLP-RAs ^6,7^, or targeted *via* multi-receptor agonists ^8–10^. However, clinical progression of Y2R agonists has been limited by poor tolerability and high rates of adverse effects^11^. Targeting nutrient- and metabolite-sensing GPCRs that drive native PYY release from enteroendocrine L-cells presents an alternative strategy for harnessing the metabolic benefits of PYY whilst potentially avoiding the onset of GI symptoms that arise from systemic activation of central circuits. PYY also promotes colonic epithelial regeneration in *in vitro* models of inflammatory bowel disease^12^, further supporting the characterisation of agonists that enhance local PYY accumulation in the gut microenvironment.

Although PYY and GLP-1 are both products of the L-cell, these hormones occupy distinct vesicular populations^13^, and certain stimuli can selectively promote the release of only one^14,15^. In contrast to the well-characterised mechanisms driving GLP-1 secretion ^16–21^, we currently have far scarcer knowledge on druggable targets controlling PYY release from the L-cell. Efforts to screen nutrient and/or pharmacological PYY secretagogues are limited due to a lack of immortalized cell lines that reliably secrete PYY^22^. Intestinal organoids - though physiologically representative - require complex differentiation protocols that yield low and variable proportions of enteroendocrine cells. Moreover, current methods for the detection of gut hormone secretion are both cost- and time-intensive. Here, we describe a novel fluorescent reporter system for measuring PYY from human L-cells, with scalable assay readouts and a functional responsiveness to bioactive nutrient stimuli.

## Results

### *In silico* construct design guides faithful posttranslational processing and targeting of SEP-PYY to secretory granules in a human enteroendocrine cell line

Our aim was to design a PYY biosensor which could faithfully mimic the biosynthesis and secretion of endogenous PYY in a human enteroendocrine cell line. The pH-sensitive variant of green fluorescent protein (GFP), super-ecliptic phluorin (SEP), was chosen as the fluorescent tag for construct design due to its high signal-to-noise ratio and appropriate pKa for reporting vesicle fusion^23^. To ensure tagging of a mature and functional PYY construct, SEP-tagged PYY (SEP-PYY) sequences included the full-length prohormone precursor of PYY (preproPYY). In the rationale design of an insertion point for the SEP tag, we considered three criteria; A) preservation of prohormone enzymatic processing sites required for correct posttranslational processing and targeting of functional PYY to secretory granules (**Figure 1a**) ^24^ B) maintenance of the conserved Y2R-binding ‘PP-fold’ domain of PYY (residues 2-8 & 12-33 of PYY1-36)^25,26^ C) integrity of the C-terminal pentapeptide (residues 32-36 of PYY1-36) is crucial for receptor binding potency^27–29^. Based on these criteria, SEP was inserted between Tyrosine1 and Proline2, and *in silico* structural predictions of the candidate sequence was determined by AlphaFold2^30^ (**Figure 1b**).

**Figure 1:**
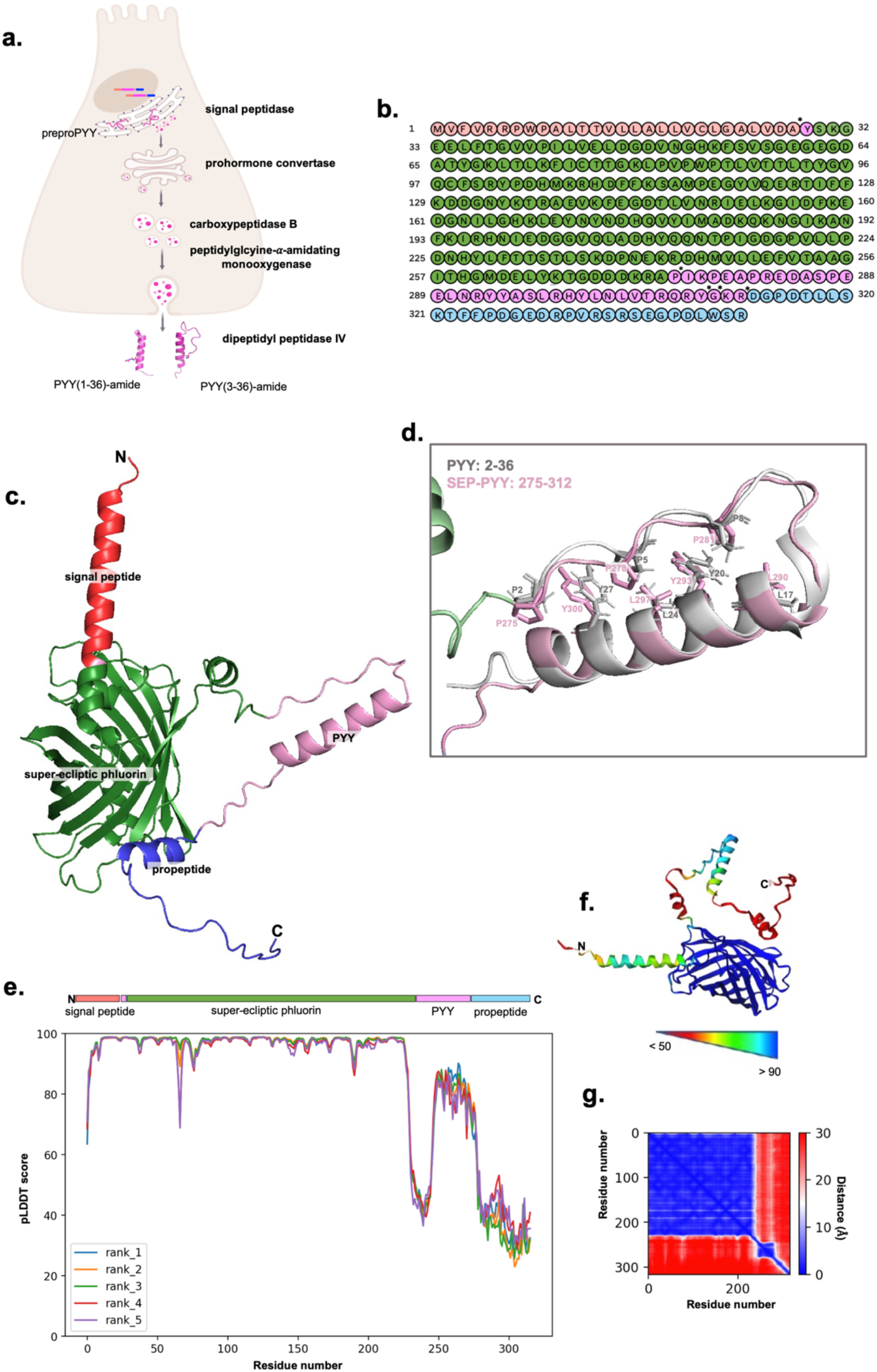
SEP-PYY construct design preserves native prohormone processing sites and structure of PYY functional domains. **a)** Schematic of post-translational processing of preproPYY in native L-cells, featuring the crystal structure of mature PYY1-36 (PDB: 7RT9). Created by BioRender. Hirdaramani. A. (2026). **b)** Amino acid sequence of SEP-PYY coloured by domain. Red: Signal Peptide, Pink: PYY1-36, Green: SEP tag, Blue: Propeptide. **c)** Highest-ranking AlphaFold2 3D structural prediction (Rank 1) of SEP-PYY. **d)** Structural overlay of the PP-fold domain of crystal structure of PYY (PDB: 7RT9) and AlphaFold2 Rank 1 prediction of SEP-PYY, coloured in grey and pink respectively, with key domain interface residues labelled. **e)** Predicted Local Distance Difference Test (pLDDT) Output for SEP-PYY. **f)** SEP-PYY Rank 1 coloured by pLDDT score. **g)** Predicted Alignment Error (PAE) output of SEP-PYY Rank 1.

Structural prediction of the top scoring model of SEP-PYY (**Figure 1c**) was superimposed with the crystal structure of human PYY1-36 (**Figure 1d**)^31^, confirming that key inter-residue distances between the N-terminal polyproline core (P2, P5 and P8 of PYY1-36; P275, P278, P281 of SEP-PYY) and the C-terminal helix of the PP-fold (L17, Y20, L24 and Y27 of PYY1-36; L290, Y293, L297 and Y300 of SEP-PYY) were within 0.6Å of each other **(Table S1)**. Per-residue and domain-level confidence was assessed using predicted local distance difference test (pLDDT) (**Figure 1e-f**) and predicted alignment error (PAE) scores respectively (**Figure 1g**). The mean pLDDT score indicated ‘High’ model confidence (averaging 85.6 across models). Residues corresponding to PP-fold (residues 276-312) achieved PAE scores lower than 10Å, which are consistent with high confidence predictions in the arrangement of this domain. Lower pLDDT and PAE scores were confined to the intrinsically disordered C-terminal tail of SEP (residues 258-275) ^32^ and the PYY propeptide region (residues 313-342). which lacks homologous structures and exhibits low evolutionary constraints **(Figure S1)**^33^. Using the SignalP server^34^, we obtained a high probability (p=0.8003) prediction of signal peptidase-mediated cleavage of SEP-PYY between positions A28 and Y29, which maps to the respective cleavage site of endogeneous preproPYY which frees the mature 1-36 form peptide **(Figure S1).**

We selected the human colonic NCI-H716 cell line, which models the nutrient-sensing receptor complement^13,35^ and GLP-1 secretion profiles^36^ of human L-cells, to express the SEP-PYY biosensor. Native PYY expression is detectable at the transcript level in this cell line, although at ∼5000 lower abundance relative to proglucagon^13,15^, and several attempts to quantify PYY secretion in this model have either failed to detect secretion^22^ or report release following chronic ligand stimulation^13^ to pre-load cellular stores^15^. SEP-PYY was transiently transfected into NCI-H716 cells, as well a non-enteroendocrine control cell line, HEK-293, and protein expression levels were assayed by flow cytometry (**Figure 2a**). SEP-PYY protein expression, quantified by Mean Fluorescence Intensity (MFI), was significantly higher in NCI-H716 cells than HEK-293 cells (**Figure 2b**). Confocal microscopy revealed a distinct vesicular expression profile of SEP-PYY n NCI-H716 cells (**Figure 2c**), consistent with post-translational processing and targeting into secretory granules. In contrast, transfection into HEK-293 cells as well as the intestinal absorptive line Caco-2, yielded diffuse cytoplasmic fluorescence (**Figure 2c**), indicating a dependence of SEP-PYY on endogeneous EEC machinery for efficient granule targeting. We next confirmed that this fluorescent signal in NCI-H716 cells was indicative of a full-length tagged SEP-PYY construct, rather than cleaved or non-specific fluorescent derivatives. Western Blotting with an anti-PYY antibody detected a 38kDa band, corresponding to the molecular weight of SEP-PYY, in lysates from transfected but not untransfected cells (**Figure 1d**). A band indicative of endogenous PYY(1-36) expression (predicted at 11kDa), was not observed in any condition, in line with the low native PYY expression in this cell line.

**Figure 2:**
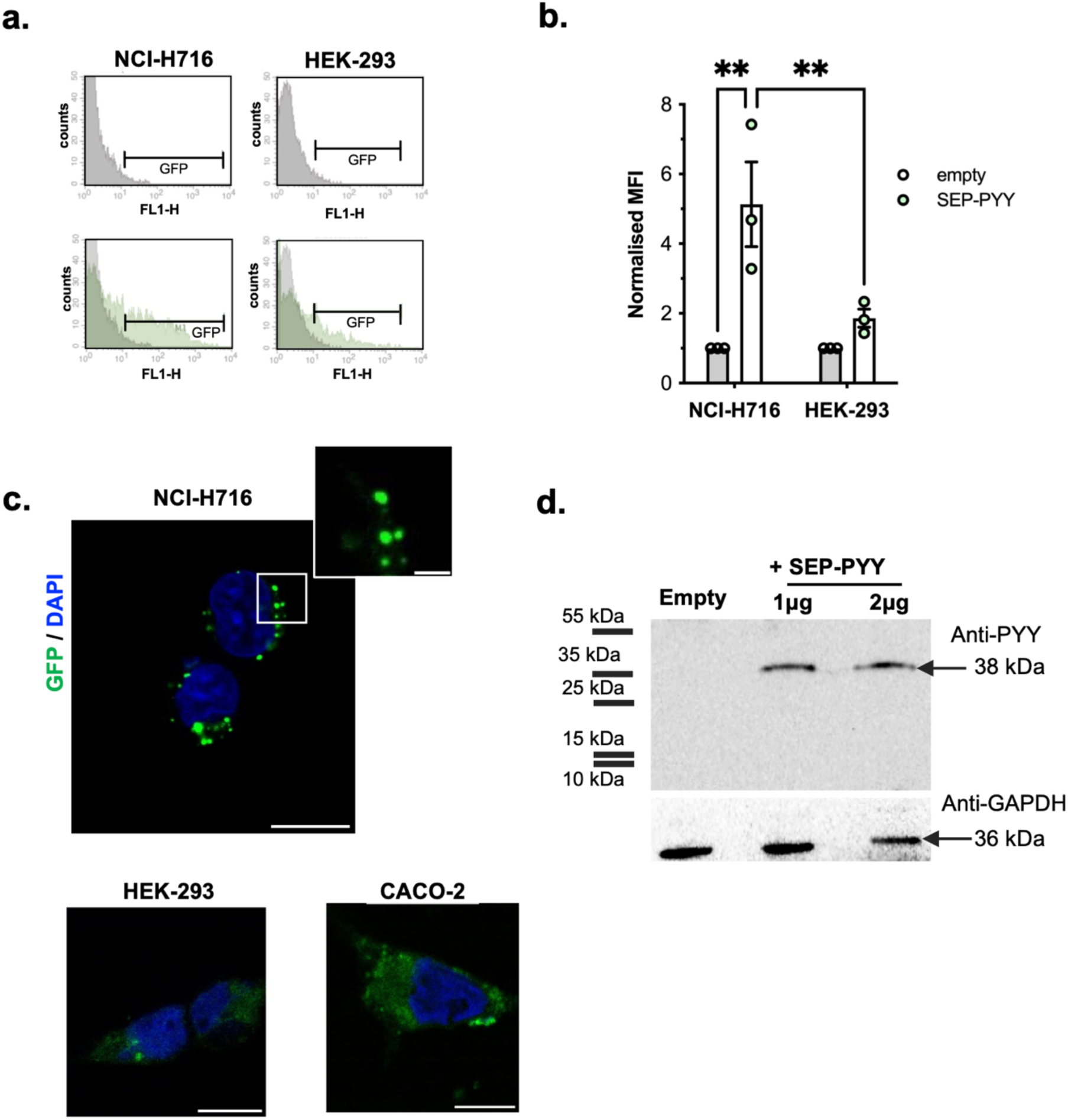
SEP-PYY is faithfully targeted to secretory granules in the human enteroendocrine NCI-H716 cell line. Protein expression levels of SEP-PYY in transfected NCI-H716 and HEK-293 cell lines were quantified by flow cytometry. **a)** Representative histograms showing GFP fluorescence (FL-H) and cell counts in untransfected (Empty) and transfected NCI-H716 and HEK-293 samples. **b)** SEP-PYY mean fluorescence intensity (MFI) normalised as fold change over untransfected cells, in NCI-H716 and HEK-293 cells. Symbols represent the mean + SEM of three individual repeats. (**p<0.01, Two-way ANOVA comparing SEP-PYY versus empty vector conditions within each cell line and SEP-PYY-transfected NCI-H716 versus HEK-293 cells**. c)** Confocal images of SEP-PYY expressed in NCI-H716, HEK-293 and Caco-2 cells. Representative images of n= 10 cells, captured across three individual repeats. Scale Bar = 10μM, Inset Scale Bar = 1μM. **d)** Western blot of NCI-H716 cell lysates probed with anti-PYY antibody and anti-GAPDH as a housekeeping control. Lysates were prepared from untransfected (Empty) cells and cells transfected with 1 μg or 2 μg SEP-PYY plasmid.

### SEP-PYY production and secretion is responsive to nutrient stimuli

We and others have previously demonstrated that the short-chain fatty acid (SCFA) butyrate robustly upregulates PYY in *in vitro* L-cell models ^13,15,37,38^. To confirm whether SEP-PYY expressing NCI-H716 cells mimic the responsiveness of endogenous PYY to upstream nutrient cues and demonstrate the utility of this biosensor across platforms, we measured SEP-PYY expression by flow cytometry, and secretion by a spectrofluorometer-based plate assay, in response to SCFA stimulation (**Figure 3a**). We quantified SEP-PYY production by flow cytometric measurements of cellular GFP fluorescence (**Figure 3b-c**). Butyrate significantly increased both the percentage of cells GFP-positive cells and mean cellular fluorescence intensity (MFI) in SEP-PYY-transfected NCI-H716 cells, consistent with the increased vesicles observed qualitatively via confocal microscopy (**Figure 3d**). In contrast, we did not detect increased SEP-PYY production following treatment with either acetate or propionate, consistent with previous findings that these SCFAs are less efficacious stimulators of native PYY production and a transcriptional/translational level than butyrate^13,15^

**Figure 3:**
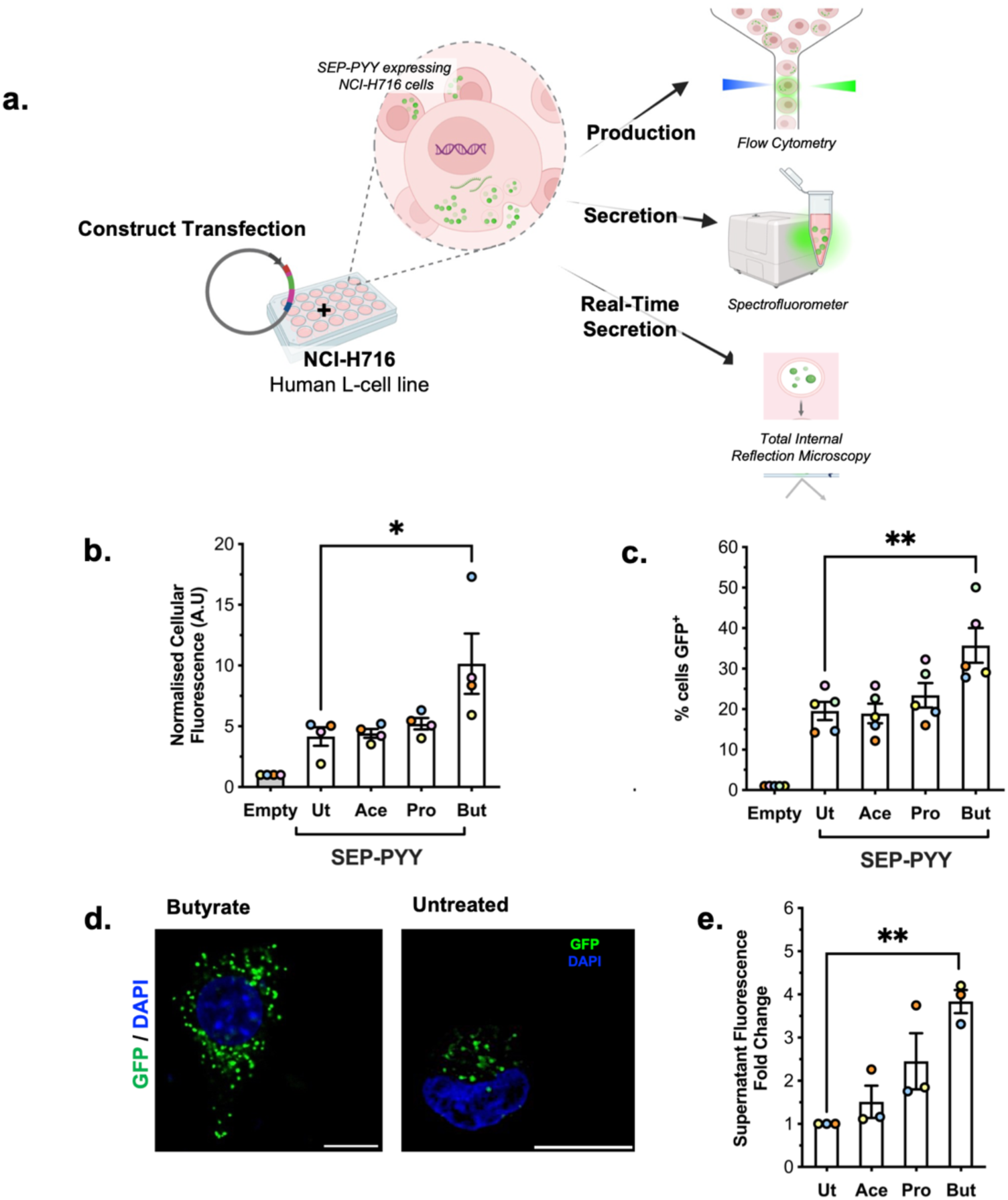
Butyrate upregulates SEP-PYY production and secretion in NCI-H716 cells. **a)** Schematic of workflows for assaying SEP-PYY production and secretion in NCI-H716 cells. Flow cytometric measurements of **b)** normalised cellular fluorescence and **c)** % cells GFP-positive in transfected NCI-H716 cells (*Empty)* and NCI-H716 cells transfected for 48 hours with SEP-PYY. Transfected cells were untreated (*Ut)* or treated with 2mM of Acetate (*Ace*), Propionate (Pro), or Butyrate (But) for 24h. Data are presented as fold change over untransfected cells. Symbols represent the mean + SEM of at least n=4 individual experimental runs. (*p<0.05; **p<0.01; One-way ANOVA with Dunnett’s post-hoc test; SEP-PYY *Ut* vs *Ace/Pro/But*). **d)** Confocal images of SEP-PYY transfected cells with Butyrate treatment, and in Untreated conditions. **e)** Supernatant fluorescence of NCI-H716 cells transfected with SEP-PYY for 48h and treated with 2mM Ace, Pro or But for 24h (or left Untreated), followed by 2hr incubation in ligand-free secretion buffer. Data are presented as fold change over untransfected cells. Symbols represent the mean + SEM of n=3 individual experimental runs. (**p<0.01; One-way ANOVA with Dunnett’s post-hoc test; SEP-PYY *Ut* vs *Ace/Pro/But*).

The detection of gut hormone secretion conventionally relies on ELISA or immunoassays, which are cost- and time-intensive respectively. We leveraged and optimised a spectrofluorometric readout for detection of SEP-PYY secretion from NCI-H716 cell, which outperforms current methods of hormone detection in speed and price. Chronic exposure of each SCFA resulted in a significant increase in SEP-PYY levels secreted into the media following butyrate treatment (**Figure 3e**).

The use of total internal reflection microscopy (TIRF) in engagement of exocytic vesicles with the plasma membrane has been valuable in investigating phasic release profiles of other endocrine peptides^39,40^, and phase-specific defects in pathophysiological conditions^41^. The pKA of SEP confers a key advantage in reporting exocytotic events; plasma membrane fusion of acidic secretory vesicles (∼pH 5.5) with the extracellular environment (∼pH 7.4) coincides with a marked increase in phluorin fluorescence (**Figure 3a**). We captured single-event SEP-PYY exocytosis in SEP-PYY-transfected NCI-H716 cells using TIRFM, acquiring videos, at 100ms intervals over 60s **(Supplementary Video 1).** Flash events appeared as prominent increases in fluorescence, exceeding a threshold defined as the mean background intensity + 2 standard deviations, consistent with established criteria for analysis of exocytosis events by fluorescence techniques^42^ (**Figure 4b**). The mean duration of individual events was 472.2 ms (**Figure 4c**), consistent with previous TIRF-M studies of peptide release ^39,41^. Using this approach, we quantified a ∼2.5 fold increase in flash event frequency with butyrate treatment, consistent with the effect of butyrate on enhanced secreted SEP-PYY fluorescence (**Figure 4d**). TIRF-M studies of insulin^41^ and GLP-1^39^ secretion have shown that flash events can arise from either pre-docked granule pools already resident at the plasma membrane or from ‘newcomer’ granules that are trafficked from deeper in the cell towards immediately before membrane fusion. In our recordings, flash events arose exclusively from newcomer granules, which appeared de novo at the plasma membrane before fusion.

**Figure 4:**
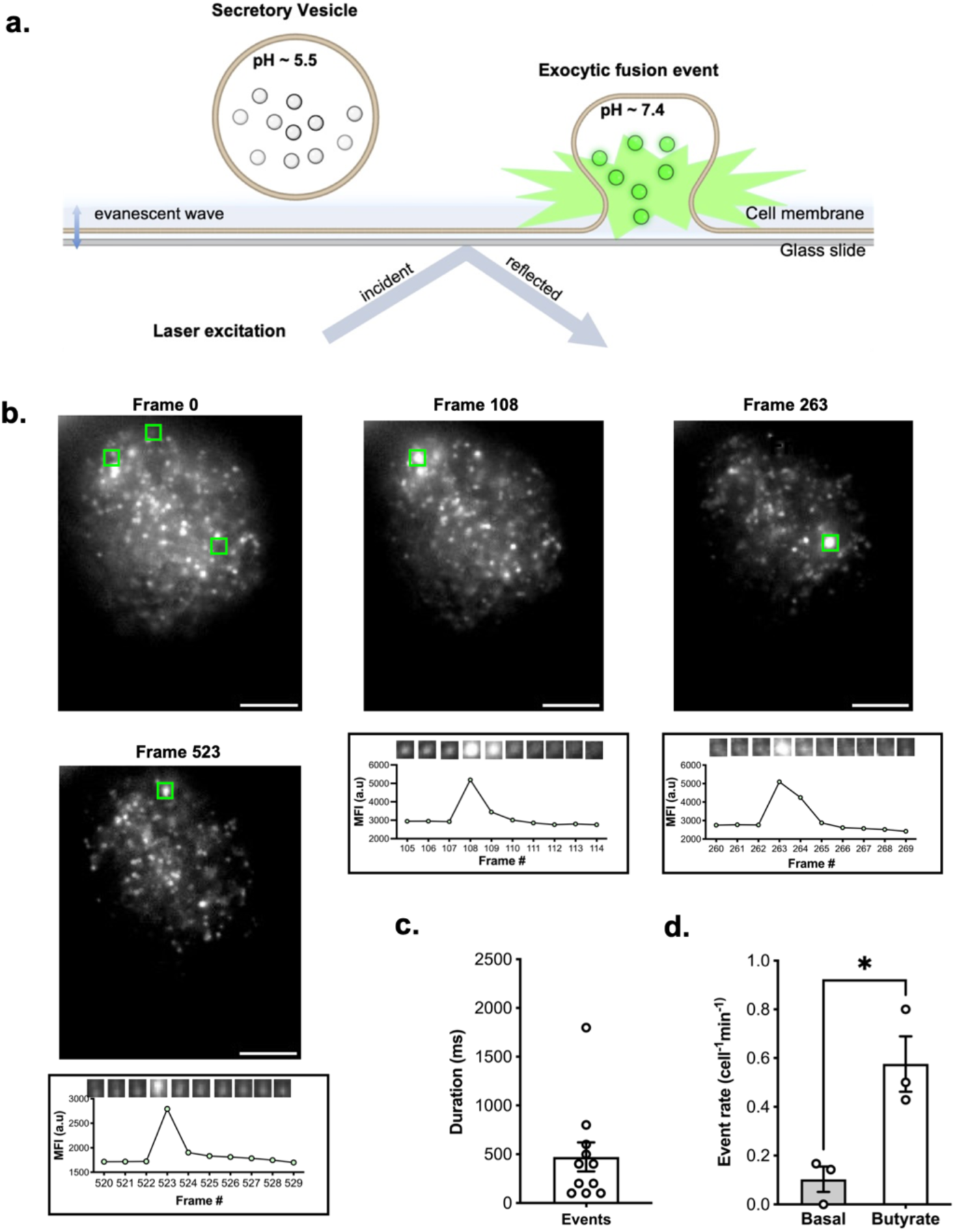
Visualising SEP-PYY secretion with single-event resolution using Total Internal Reflection Fluorescence Microscopy (TIRF-M). **a)** Schematic of a characteristic single SEP-PYY vesicle exocytic fusion event at the plasma membrane captured by TIRF-M. **b)** Representative image sequence of a SEP-PYY-transfected NCI-H716 cell, pre-treated with 2mM But for 24h, captured by TIRF-M resolved at 100ms intervals, for 60 seconds. Green borders indicate regions of interest (ROI) for flash events. Associated traces show the mean fluorescence intensity (MFI) of each flash event sampled at100ms intervals, alongside zoomed views of each ROI in the indicated frame range. **c)** Mean flash event duration in SEP-PYY transfected NCI-H716 cells, captured in n=11 cells, across n=3 independent experiments. Error bar represents SEM. **d)** Quantification of flash events in SEP-PYY transfected cells pre-treated with butyrate, or without treatment (Basal). n=22-24 cells were imaged per condition, across n=3 independent experiments. Symbols represent the mean ± SEM of each repeat. (**p<0.01, t-test; *Ut* vs *But*).

### SEP-PYY can be used to screen nutrient and pharmacological agonists of L-cell GPCRs for their ability to act as acute secretagogues

A diverse complement of nutrient-sensing GPCRs drive enterohormone release in native L-cells, presenting multiple targetable pathways for synergistic stimulation of secretion^43^. We tested the responsiveness of the SEP-PYY biosensor to acute stimulation with ligands of GPCRs which are endogenously expressed in L-cells, and have previously been implicated in driving GLP-1 release from NCI-H716 cells^13,21,44–46^ (Figure 5) ^17,47–50^. SEP-PYY-transfected cells were treated for 2 hours with stimuli of either free fatty acid receptor 2 (FFAR2), free fatty acid receptor 4 (FFAR4), bile acid receptor (gpbar1), GPR119 or calcium-sensing receptor (CaSR). We detected a significant increase in fluorescence with either the FFAR2 agonist propionate or the GPR119 lipid agonist oleoylethanolamide (OEA) (Figure 5), indicating that although these receptors are all known to induce GLP-1 secretion, they may differ in their ability to induce secretion of PYY.

**Figure 5:**
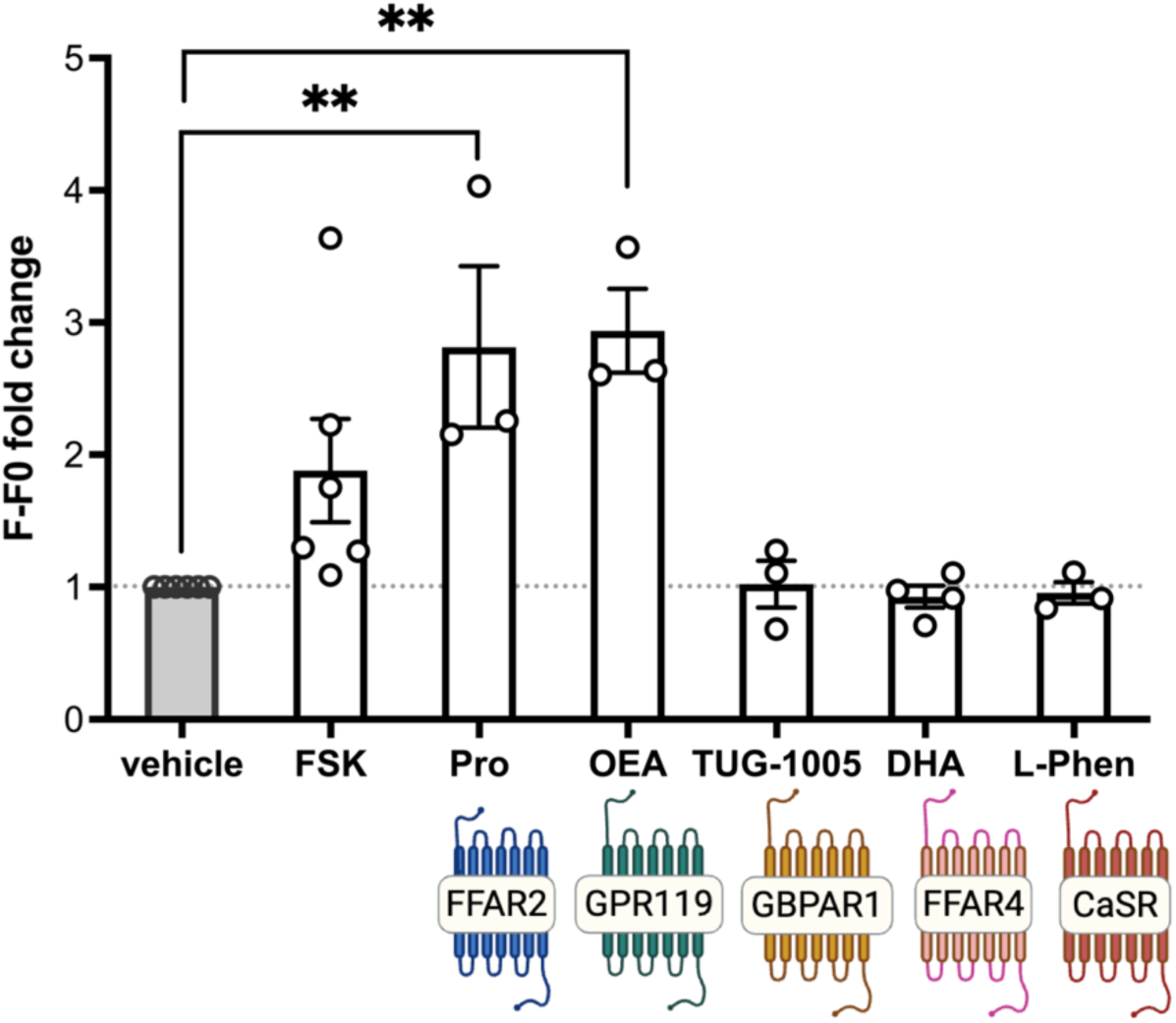
SEP-PYY secretion is differentially regulated by agonists of nutrient-sensing GPCRs in NCI-H716 cells. Supernatant fluorescence of NCI-H716 cells transfected with SEP-PYY for 48h and treated for 2 hours with 10uM Forskolin (FSK), 10mM Propionate (Pro), 10uM oleoylethanolamide (OE), 10uM TUG-1005, 100uM docasohexanoic acid (DHA) and 30mM L-phenylalanine (L-Phen) in ligand-free secretion buffer. Data are presented as background-corrected fold change over untransfected cells. Symbols represent the mean + SEM of at least n=3 individual experimental runs. **p<0.01; one-way ANOVA with Dunnett’s post-hoc correction.

## Discussion

Here, we establish a scalable *in vitro* system for investigating real-time, stimulus-evoked PYY profiles in human EECs using a SEP-tagged PYY biosensor. We show that the reporter is dependent on native enteroendocrine machinery for processing and targeting into secretory granules, and stimulated by canonical nutrient stimuli, enabling inference of physiologically-relevant hormone output. Our optimised workflows provide an approximate fivefold reduction in assay time (∼20 min readout) and ninefold reduction in per-sample cost (∼1USD) relative to ELISA-based detection, which maintaining sensitivity for both biosynthesis and secretion. compared to ELISA-based detection methods. This system could also be readily adapted to more complex epithelial models such as intestinal organoids, and re-engineered to interrogate alternative hormone cargos.

EECs express a wide repertoire of nutrient-sensing GPCRs that represent attractive opportunities for selectively modulating gut hormone release^1^. Previous work^13,51^, including our own, show that GLP-1 and PYY exist in distinct vesicle pools in enteroendocrine L-cells, and exhibit non-identical stimulus-secretion coupling. We therefore screened SEP-PYY expressing cells with a panel of GPCR agonists previously implicated in driving GLP-1 release from L-cells^13,21,44–46^, we found that only acute activation of FFAR2 and GPR119 elicited robust PYY release, revealing a degree of input selectivity via mechanisms that are likely beyond heterotrimeric G protein coupling profile alone, but could be exploited to fine-tune enterohormone profiles while limiting off-target central effects. Moreover, leveraging the rich signalling diversity of the GPCR superfamily – including subcellular compartmentalisation^13,52,53^, receptor cross-talk^54,55^, and ligand-biased signalling^56^ – offers superior routes to precision control of gut hormone secretion.

We demonstrate the use of this platform paired with TIRF-M in reporting live single-event hormone exocytosis. Moreover, our live-cell imaging uncovered distinct granule dynamics for PYY dynamics compared to GLP-1. Previous imaging studies have shown biphasic exocytosis of glucose-stimulated GLP-1 secretion which originates from pre-docked granule pools, followed by newcomers^39^. In contrast, SEP-PYY flash events in our TIRF-M recordings arose exclusively from new vesicle events, with no pre-docked granule pools observed. This further supports the concept that GLP-1 and PYY secretion are regulated in distinct manners in addition to perhaps distinct secretory pools. Current understanding of PYY secretion mechanisms are limited, thus, coupling this biosensor with live advanced imaging modalities will enable dissection of regulatory mechanisms governing PYY production, granule trafficking and exocytosis, and may help identify potential phase-specific defects in conditions associated with lowered PYY release ^57,58^.

Although we have demonstrated that SEP-PYY replicates enteroendocrine-restricted faithful expression and largely recapitulates the functional behaviours of endogenous PYY, we acknowledge that a biosensor requires overexpression of an exogeneous construct, and although a range of ligands were assessed in this study, it may distort native secretory dynamics, such as with vesicle targeting and release, thereby skewing quantitative measures of basal and stimulus-evoked output. Future studies could employ gene-editing approaches to label endogenous PYY, although this approach in turn has its own limitations in terms of off target effects.

Despite the heightened popularity of gut hormone receptor analogues in obesity, diabetes and beyond, these agents have important limitations. Gastrointestinal disturbances are common, and their benefits require long-term usage because weight gain is frequent following discontinuation^59^. In contrast, bariatric surgery achieves durable weight loss, which has been associated with postoperative elevation of both PYY & GLP-1 secretion in these patients^60^, implicating enhanced endogenous gut hormone secretion in mediating favourable metabolic outcomes. In order to mimic the sustained benefits of bariatric surgery, next-generation therapeutics could converge on identifying ligands to promote enterohormone secretion from native EECs, rather than relying on exogeneous analogues. Creation of tools, such as the PYY biosensor in this study, could aid such efforts in addition to providing further understanding of the triggers, mechanisms and extracellular environmental requirements that drive PYY secretion in the gut, both physiologically and in disease.

## Author Contributions

*Conceptualization & Experimental Investigation*: A.H.; *Resources*: A.C.H. and G.F.; *Writing:* A.H.; *Editing,* A.C.H.; *Visualization,* A.H.; *Supervision*, A.C.H. and G.F.; *Funding Acquisition:* all authors.

## Methods

### Cell Culture

NCI-H716 cells (ATCC, CCL-251) were maintained in suspension in RPMI-1640 (Sigma-Aldrich) containing 2g/L D-glucose and 2mM glutamine supplemented with 10% (v/v) fetal bovine serum (Merck) and 100U/ml penicillin/streptomycin (Sigma-Aldrich) (complete RPMI). Two days before experiments, cells were seeded onto culture plates were pre-coated with Cultrex Basement Membrane Extract (170μl/cm^2^) (R&D Systems) diluted to 0.5mg/ml. HEK-293 (ATCC, CRL-1573) and Caco-2 (ATCC, CRL-2102) cells were grown as an adherent monolayer in T-75 flasks in DMEM containing 4.5g/L D-glucose and 4mM glutamine supplemented with 10% (v/v) fetal bovine serum (FBS) and 100U/ml penicillin/streptomycin (+/+ DMEM).

### Construct Design & Molecular Cloning

Custom gene fragments containing the human PYY sequence **(ID)** tagged with super-ecliptic pHluorin **(ID)** were synthesised *de novo* by Twist Bioscience. Amino acid sequences of SEP-PYY constructs were input to AlphaFold2 (v2.) for structural predictions. Five model predictions were generated, with 3 recycles for structural refinement. Model confidence was assessed using pLDDT scores and PAE matrices. Structural preservation of functional domains between endogenous human PYY and the highest confidence (rank 1) SEP-PYY models was evaluated by measurement of inter-residue distances on Pymol (v). Signal peptidase I cleavage sites were predicted using SignalP5.0 software. Gene fragments were subcloned into pcDNA 3.1+/+ mammalian expression vector using *XbaI* and *AfeI* restriction enzyme sites.

### Transfections

SEP-PYY constructs were expressed in NCI-H716 cells by transient reverse transfection using Lipofectamine 2000 reagent. SEP-PYY-Lipofectamine 2000 mixes were prepared (1ug DNA:5ul Lipofectamine/well) and transferred to 6 well plates pre-coated with Matrigel (0.5mg/ml), and cell suspensions were then added immediately. For transfection of SEP-PYY into the HEK-293 cell line, cells were plated and grown to 75-85% confluency prior to transfection. SEP-PYY-Lipofectamine 2000 mixes were prepared as above and added dropwise to cell culture media. Media was changed 16-20 h after transfections, and fresh media contained respective SCFAs in the case of chronic SCFA treatments. Cells were grown for an additional 24h, till 70-80% confluent. Reagent volumes and cell number were scaled down based on the surface area of culture plates used for used for specific assay formats.

### Flow Cytometry

SEP-PYY production was assessed by flow cytometry. Cells were washed twice with cold phosphate-buffered saline (PBS) with Ca2+ (PBS-Ca2+) and harvested in FACs buffer (PBS supplemented with 2% FBS). Samples were centrifuged for 5 min at *1118 x g* at 4°C, resuspended in FACs buffer and transferred to polypropylene round bottom FACs tubes. Data was acquired a FACS Calibur Flow cytometer. The parameters of the gating region of interest were determined using readings with untransfected cells.

### Fluorometer-based Secretion Assay

SEP-PYY secretion was assessed by measurement of fluorescence with a microplate reader-based protocol. On the day of the assay, cells were washed once with PBS and incubated with respective ligands diluted in Krebs buffer (supplemented with 0.2% BSA) for 2h at 37C. Secretion buffer was then collected, immediately buffered with 15mM HEPES (pH 7.4) and spun down for 10 min at *800 x g* at room temperature. The solution was then transferred to 96 well clear bottom black walled microplates (Corning) and fluorescence was measured on a LUMIstar OPTIMA microplate reader with optical filters for excitation and emission set to 485nm and 520nm, respectively, using bottom optics. Wells were scanned using orbital averaging of four measurements per well, and detector gain was set to 70% to avoid saturation. Raw fluorescent values were background-subtracted (using no-cell treatment controls) and normalised to signals obtained from untransfected cells in parallel.

### Confocal microscopy

For confocal microscopy, glass coverslips (13mm) were added to the bottom of wells prior to Matrigel coating, reverse transfections and cell seeding. On the day of the assay, cells were washed with cold PBS-Ca2+ prior to fixation with 4% paraformaldehyde for 20 minutes at room temperature. Cells were washed and coverslips were mounted using Fluoromount G onto clear glass slides. Slides were imaged on a Stellaris 8 Inverted confocal microscope (Leica) using the LasX software.

### TIRF microscopy

Total internal reflection fluorescence microscopy was conducted on a NanoImager S from ONI (Oxford Nanoimaging; ONI) with cells plated on ibidi μ-Slide 8 well glass bottom chambers. 30 minutes before imaging, cell media was changed to Krebs buffer supplemented with 0.2% BSA, containing respective ligands. Frames were captured on a close-loop piezo stage with a 100x NA 1.4 oil immersion objective and SCmos camera with excitation 405nm, 488nm, 561nm and 640nm lasers, and temperature at the nanoimager was controlled at 37C. Samples were brought into TIRF mode by adjustment of the angle of incidence of laser excitation. Frames were captured every 100ms (10 frames per second) for 5 minutes per sample.

### Western Blotting

Transfected NCI-H716 cells were harvested and lysed with RIPA lysis buffer. Cell lysates were mixed with 4x Laemli Sample buffer with 5% 2-mercaptoethanol and heated at 100°C for 10 min. Samples were resolved by SDS-page at 150V for 2 h, and proteins were transferred to a 0.22 μM nitrocellulose membranes at 100V for 90 min at 4°C. Membranes were blocked in 5% non-fat dry milk in TBS-Tween (blocking buffer). After washing, membranes were incubated with an anti-PYY antibody (rabbit polyclonal, bioRT, orb579781) at a 1:250 dilution in blocking buffer, overnight at 4°C. Membranes were washed and then incubated with HRP-conjugated anti-rabbit secondary antibody (Invitrogen, 31460) at 1:10,000 dilution in blocking buffer for 1h at room temperature. Membranes were developed with Immobilon Forte Western HRP Substrate, and imaged using ImageQuant sofrware. Band density was quantified normalised to GAPDH.

### Statistical Analysis

All statistical analyses were carried out as per indicated in figure legends on GraphPad Prism. “Unpaired t-tests were used to compare two groups. One-way ANOVA with Dunnett’s post hoc test was used for comparisons of multiple groups to the untreated control. Two-way ANOVA was used for experiments involving multiple groups and conditions.

### Materials Availability

SEP-PYY construct is available from the lead contact under a materials transfer agreement with Imperial College London.

---

**Supplementary Table 1.** Measured distances in angstroms between key structure-function residue pairs of NMR-resolved structure of PYY (PDB: 2DEZ) and AlphaFold2 3D prediction of SEPPYY visualised on Pymol.

| PYY1-36 |  | SEP-PYY |  |
| --- | --- | --- | --- |
| Residue pair | Cα-Cα (Å) | Residue pair | Cα-Cα (Å) |
| P2-Y27 | 5.9 | P275-Y300 | 5.9 |
| P5-Y27 | 7.7 | P278-Y300 | 7.5 |
| P5-L24 | 5.7 | P278-L297 | 5.8 |
| P5-Y20 | 6.0 | P278-Y293 | 6.6 |
| P8-Y20 | 6.6 | P281-Y293 | 7.1 |
| P8-L17 | 6.3 | P281-L290 | 6.4 |

**Supplementary Figure 1.**
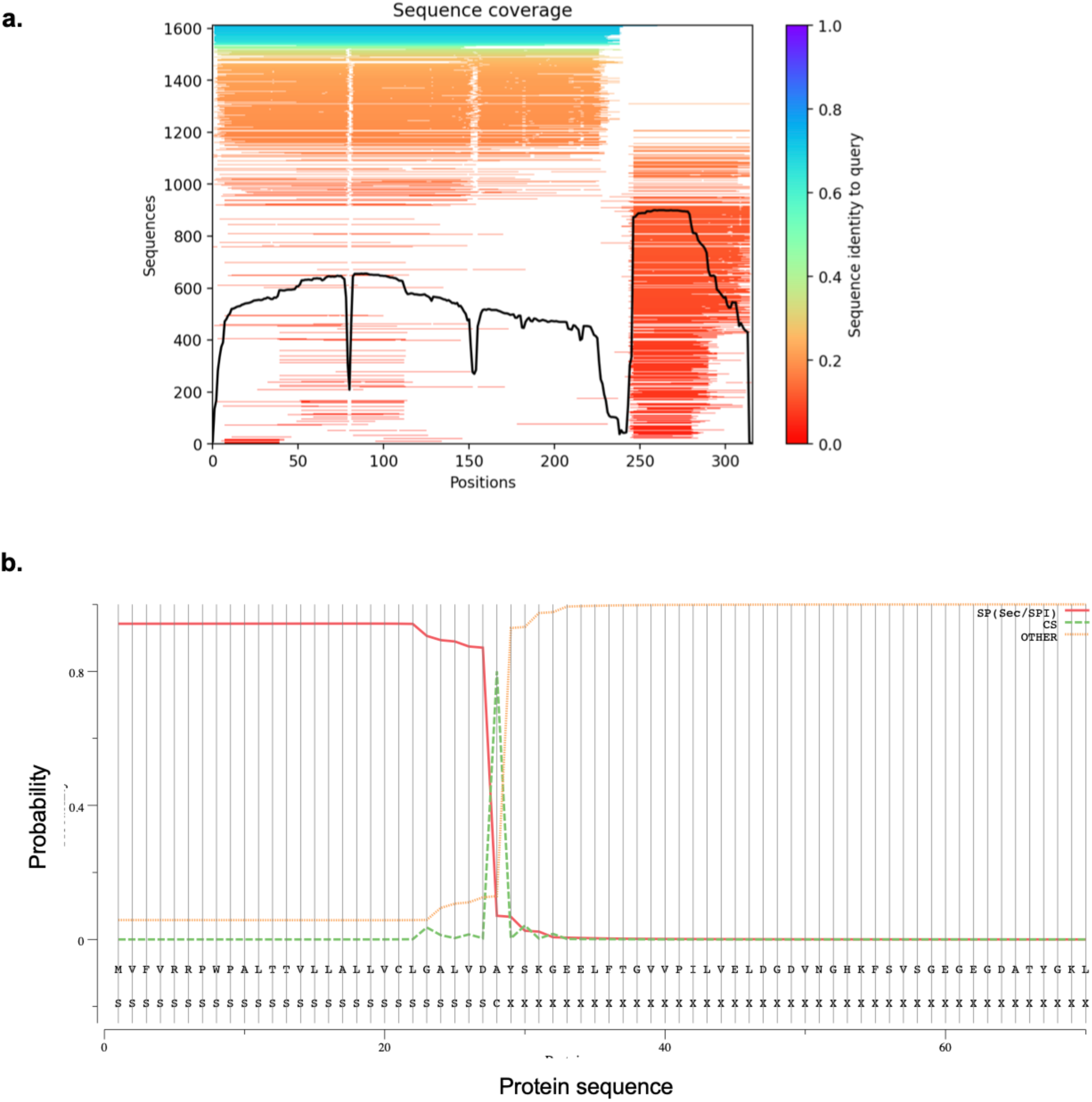
a) AlphaFold2 MSA sequence coverage plot of SEP-PYY. B) b) SignalP 5.0 predicted probability of signal peptide sequence (red solid line), cleavage site (dashed green line) or other (dotted orange line) along residues 0-70 of SEP-PYY.

