## Supplementary figures and images for "A fluorescent reporter system for scalable and live detection of PYY production from enteroendocrine cells with single-event resolution"

### Supplementary Figure 1

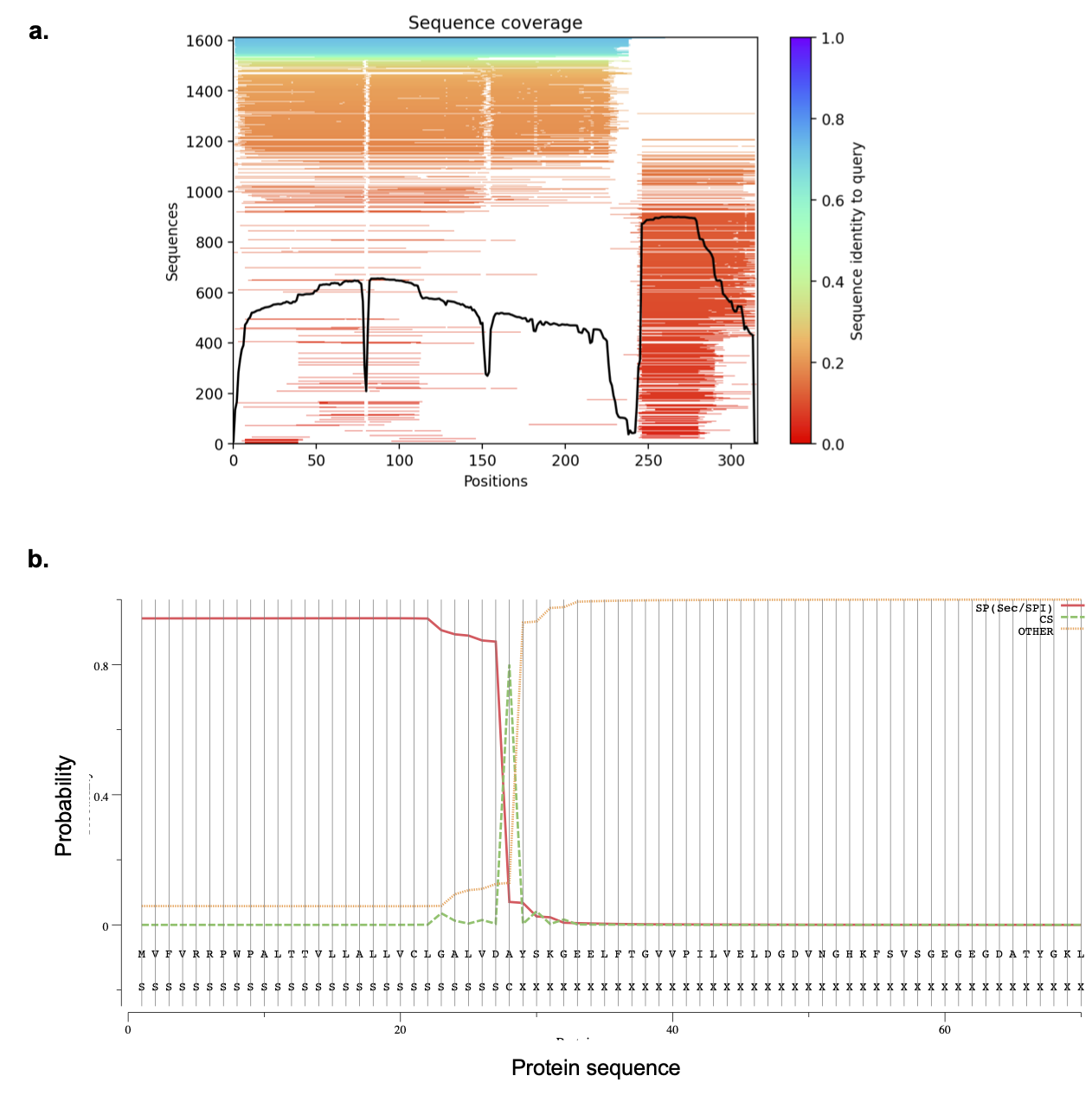

### Supplementary Video 1

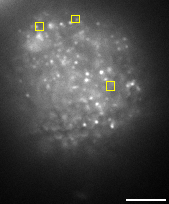
